# A study on the protective effects of CpG ODN-induced mucosal immunity against lung injury in a mouse ARDS model

**DOI:** 10.1101/471896

**Authors:** Guan Wang, Zong-Jian Liu, Xuan Liu, Feng-Ge Liu, Yan Li, Yi-Bing Weng, Jian-Xin Zhou

**Affiliations:** Department of Critical Care Medicine, Beijing Tian Tan Hospital, Capital Medical University, Beijing 100050, China; Department of Critical Care Medicine, Beijing Luhe Hospital, Capital Medical University, Beijing 101100, China; Center Laboratory, Beijing Luhe Hospital, Capital Medical University, Beijing 101100, China

**Keywords:** animal model of human disease, acute respiratory distress syndrome, CpG ODN; mucosal immunity, immunity routes, protection

## Abstract

This study aims to determine the feasibility of using oligodeoxynucleotides with unmethylated cytosine-guanine dinucleotide sequences (CpG ODN) as an immunity protection strategy for a mouse model of acute respiratory distress syndrome (ARDS). This is a prospective laboratory animal investigation. 20-week old BALB/c mice in Animal research laboratory were randomized into groups. An ARDS model was induced in mice using lipopolysaccharides. CpG ODN was intranasally and transrectally immunized before or after the 3^rd^ and 7^th^ day of establishing the ARDS model. Mice were euthanized on day seven after the 2^nd^ immunization. Then, retroorbital bleeding was carried, out and the chest was rapidly opened to collect the trachea and tissues from both lungs for testing. CpG ODN significantly improved the pathologic impairment in mice lung, especially after the intranasal administration of 50 ug. This resulted in the least severe lung tissue injury. Furthermore, IL-6 and IL-8 concentrations were lower, which was second to mice treated with the rectal administration of 20 μg CpG ODN. In contrast, the nasal and rectal administration of CpG ODN in BALB/c mice before LPS immunization did not appear to exhibit any significant protective effects. In conclusion, the intranasal administration of CpG ODN may be is a potential treatment approach to ARDS. More studies are needed to further determine the protective mechanism of CpG ODN.

## Introduction

Acute respiratory distress syndrome (ARDS) is a commonly observed critical illness with a rapid onset and high mortality rate (Rubenfeld, et al., 2005). Its main pathophysiological characteristics are increased pulmonary capillary permeability, pulmonary edema, pulmonary hyaline membrane changes, and pulmonary fibrosis (ARDS Definition Task Force, et al., 2012). The pathogenesis of ARDS is presently not completely clear (Han S and Mallampalli RK, 2015). At present, drug treatments are used to control this injury, but do not produce ideal outcomes (Anzueto A, et al., 1996; Dellinger RP, et al., 1998; Iwata K, et al., 2010; National Heart, Lung, and Blood Institute Acute Respiratory Distress Syndrome (ARDS) Clinical Trials Network, et al., 2011; National Heart, Lung, and Blood Institute ARDS Clinical Trials Network, et al., 2014; Rice TW, et al., 2011; Paine R 3rd, et al., 2012; Peter JV, et al., 2008), while some biological therapies are presently in the experimental or clinical trial phase (Boyle AJ, et al., 2014), and the efficacy remains to be determined. Considering that ARDS inflammatory responses mainly occur on the mucosal surfaces of the lungs (alveoli, and pulmonary capillary membranes), limiting or eliminating these responses may decrease the severity of lung tissue injury and shorten the duration of inflammatory responses, thereby preventing the occurrence or development of ARDS. Therefore, the investigators considered the possibility of whether mucosal immunity can play a limiting or eliminating role in preventing the occurrence or development of ARDS.

Mucosal immunity refers to the immune system present on the mucosa itself, which is different from systemic immunity. The route by which mucosal immunity is produced can be summarized as follows: After M cells on Peyer’s patches in the mucosal surfaces have engulfed the antigens, these migrate to the submucosa and deliver unmodified antigen to dendritic cells (DCs). Then, these DCs modify the antigens, and present these to T and B lymphocytes that have migrated from the blood vessels. After these B lymphocytes are sensitized by antigens, these differentiate into plasma cells and secrete large amounts of monomeric immunoglobulin A (IgA) to the mucosal surface. This monomeric IgA forms polymeric IgA through the binding of J chains to neutralize more pathogens. In contrast, sensitized T lymphocytes produce interleukin (IL)-2, IL-4, IL-5, IL-6, IL-8 and IL-10, and other cytokines, which can all stimulate B-cells to produce IgA. In addition, activated B cells may induce mucosal immunity in multiple distal sites through their homing route. Therefore, DC-activated B-cells and T-cells, and IgA all constitute the mucosal defense system, and play roles in neutralizing and eliminating pathogens (Dennehy PH, 2005; Featherstone C, 1997; Katial RK, et al., 2002; Lundholm P, 2002; Maxim RB, 2002). The activation pathways commonly used in mucosal immunity involve the nasal mucosa or rectal mucosa. Recent studies have shown that oligodeoxynucleotides with unmethylated cytosine-guanine dinucleotide sequences (CpG ODN) are good protective agents for mucosal immunity, and can regulate coagulation defects caused by endothelial injury (El Kebir D, et al., 2015; Li Lina, 2010; Li Xuan, 2014; Reiss LK, et al., 2012).

In summary, the nasal and rectal administration of CpG ODN was carried out in a mouse ARDS model, in order to evaluate the degree of lung injury, and understand whether the induction of mucosal immunity has a protective effect on lung injury in ARDS.

## Materials and Methods

### Animals

The study is a mouse’ animal experiment that was approved by the center of experiment animal ethics committee, Capital Medical University (Approval No.: AEEI-2017-138). CpG ODN, specific pathogen free (SPF)-grade 20-week old BALB/c mice (approval number: SCXK(Jing)2016-0011), weighing 24-26 grams with equal number of males and females, were randomized into groups. Then, CpG ODN (TCCA<u>TGACGTT</u>CC<u>TGACGTT)</u> was synthesized, and the entire sequence was modified with phosphonothioate, and phosphorylated at both ends (Sangon, Shanghai, China). The lipopolysaccharide (LPS) was purchased from Sigma Inc. (Sigma, Missouri, United States).

### Grouping

Forty-five BALB/C mice were randomized into nine experimental groups (A–J). Before the start of the experiment, blood was collected from the tail vein as a blank control (Table 1).

**Table 1.**
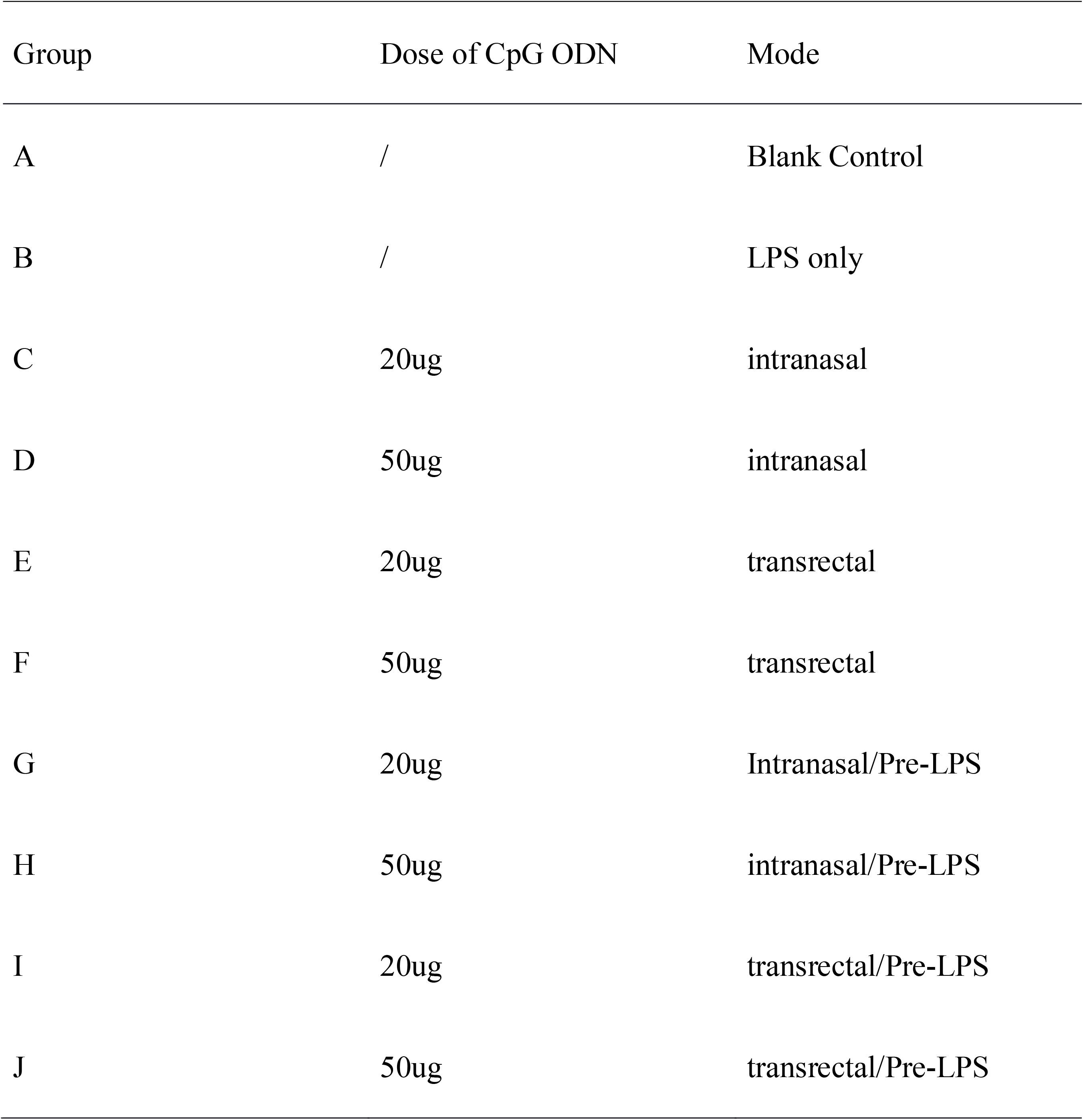
each group immunization mode on experiment

### Generation of the ARDS model (Aeffner F, et al., 2015)

Mice in groups B–F were first used to generate the ARDS models. After intraperitoneal injection of 50 mg/kg pentobarbital sodium for anesthesia, a 1.5-cm incision was made at 2 cm above the sternum to expose the trachea. Then, a microinjector was used for intratracheal instillation with 70 μg/kg of LPS to cause injury and generate the model. An absorbable suture was used, and the incision was rapidly sutured before iodophor disinfection. Mice in groups G–J initially received CpG ODN before ARDS was induced (see below). Group A was assigned as the blank control group, in which only blood and tissue samples were collected for testing, and no treatment was carried out.

### Immunization method

On day 3 and 7 after the model was generated, mice in groups C-F were given CpG ODN at 20 μg (intranasally), 50 μg (intranasally), 20 μg (transrectally), and 50 μg (transrectally), respectively. Mice in group B were only used for model generation, and CpG ODN was not administered. Mice in groups G-J received CpG ODN at 20 μg (intranasally), 50 μg (intranasally), 20 μg (transrectally), and 50 μg (transrectally), respectively, on day 1 and 3. Then, the ARDS models were generated on day seven after the 2^nd^ immunization, following the above method. Mice were euthanized on day seven after the 2^nd^ immunization in groups C–F, at day seven after model generation in groups G–J, and on the same day in group B, before retroorbital bleeding was carried out and the chest was rapidly opened to collect the trachea and tissues in both lungs. The right lung was rapidly placed in a fixative, and sent to the Pathology Department. The left lung and tracheal tissues were rapidly placed in a 4°C solution with trypsin inhibitors, and ultrasonicated. Subsequently, the homogenates were centrifuged in a 4°C low-temperature centrifuge at 12,000 rpm for 30 minutes. Then, the clear supernatant was collected and stored at –80°C for future tests. Each blood sample was centrifuged at 3,000 rpm for 10 minutes, and the clear supernatant was collected and stored at −20°C.

### Enzyme-linked immunosorbent assay (ELISA)

The levels of the inflammatory mediators (Il-6 and IL-8) were tested in the serum and tissue supernatants. IL-6 and IL-8 ELISA test kits (AMEKO ELISA Kit, Shanghai, China) were used to quantify the IL-6 and IL-8 concentrations in each sample.

### Pathological examination of lung tissues

Hematoxylin and eosin (H&E) staining of lung tissues and lung injury evaluation scores: Conventional paraffin immersion, embedding, sectioning and H&E staining were carried out before the pathological sections were observed under an ordinary microscope (magnification, ×200). A pathologist was responsible for evaluating the lung injury under a magnification of 400×. The modified grading scheme of Matute-Bello was used to evaluate the severity of lung injury, according to neutrophils in the alveolar space (S), neutrophils in the interstitial space (T), the proteinaceous debris that filled the airspaces (U), and alveolar septal thickening (V) (Aeffner F, et al., 2015). The total lung injury score was the summation of the aforementioned items. At least 20 high-powered fields (400×) were observed for each sample, and the mean value was taken (Table 2).

**Table 2.** Modified grading scheme of Matute-Bello(Chen J, et al., 2016)

| Parameter | Score per field |  |  |
| --- | --- | --- | --- |
|  | 0 | 1 | 2 |
| Neutrophils in the alveolar space | None | 1-5 | >5 |
| Neutrophils in the interstitial space | None | 1-5 | >5 |
| Proteinaceous debris filling the airspaces | None | 1 | >1 |
| Alveolar septal thickening | <2× | 2×-4× | >4× |
| Total score= [(20×S) +(14×T) + (7×U) + (2 ×V)]/number of high power fields counted ×72) |  |  |  |

### Statistical methods

Data were entered in an Excel file. Data organization and statistical analysis was carried out using Microsoft Excel (Microsoft Ltd. USA), SAS 9.4 (SAS Institute Inc., Cary, USA), and other software. Quantitative data that conforms to the normal distribution were presented as mean ± standard deviation. Intergroup comparisons were carried out using one-way analysis of variance (ANOVA). Multivariate analysis was carried out using logistic regression. A *P*-value <0.05 was considered statistically significant.

## Results

### Immunization completion status in mouse in groups A~J

In group A, all mice survived up to the end-point. In group B, four mice survived up to the end-point (B3 died on the day after the 2^nd^ LPS immunization). In group C, five mice survived up to the end-point. In group D, four mice survived up to the end-point (D2 died on the day after the 1^st^ LPS immunization). In group E, four mice survived up to the end-point (E2 died on the day after the 1^st^ LPS immunization). In group F, five mice survived up to the end-point. In group G, three mice survived up to the end-point (G1 died on the day after the 1^st^ nasal administration of CpG ODN and G3 died on the day after the 1^st^ LPS immunization). In group H, three mice survived up to the end-point (H3 and H5 died on the day after the 2^nd^ rectal administration of CpG ODN). In group I, four mice survived up to the end-point (I2 died within 10 minutes of the 1^st^ LPS immunization). In group J, four mice survived up to the end-point (J4 died on the day after the 1^st^ LPS immunization).

### Mouse ARDS model generation status

The gross appearance of the isolated lung tissues revealed congestion and edema (Fig. 1A). The pathological sections revealed that various experimental groups exhibited varying degrees of alveolar edema, the infiltration of inflammatory cells, bleeding, and atelectasis (Fig. 1B), which indicated that the ARDS models were successfully generated (Aeffner F, et al., 2015).

**Figure 1.**
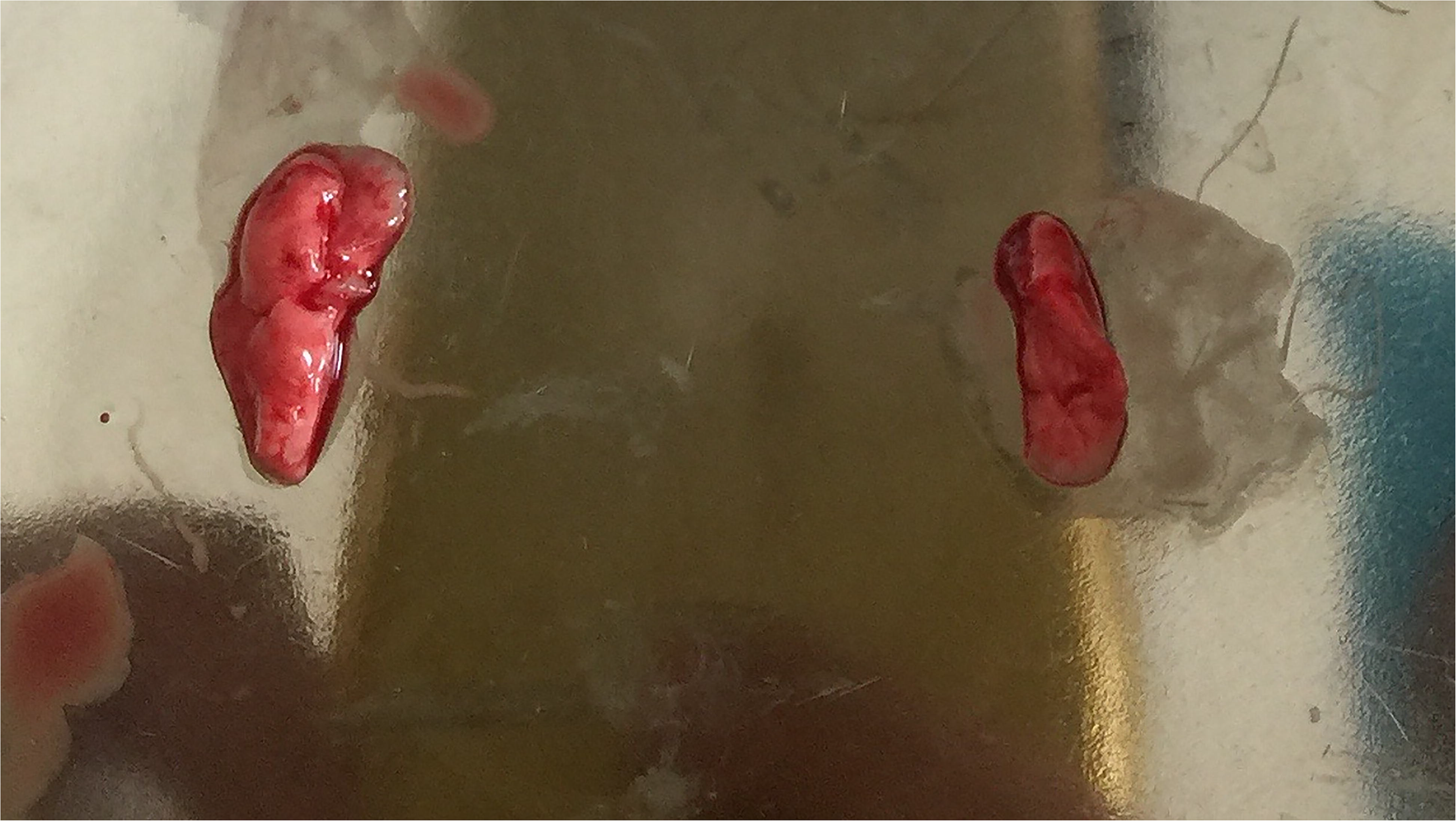
Gross appearance of lung tissue and pathology. Fig. 1A: Gross appearance of both lungs isolated from BALB/c mice after the experiment Fig. 1B: Alveolar and interstitial edema, congestion, and neutrophil infiltration in the right lung of BALB/c mice; pink protein-like substance deposition in the alveolar cavity can be seen.

### Mouse lung histopathological examination results

Group A was assigned as the blank control group. In the pathological sections (100×, 200×, and 400×), the lung tissue structure was clear, the alveolar cavity was clean, the alveolar septum was thin, and there was an absence of interstitial congestion, an absence of neutrophil elevation and migration, and an absence of protein-like substance deposition. Group B was assigned as the LPS-only group. From the pathological sections, it could be observed that the normal structure of lung tissues nearly disappeared, the alveolar cavity was significantly reduced or filled with pink protein-like substances, the alveolar septum was significantly thickened (thickness increased by >4 times compared with group A), and there was interstitial congestion and large amounts of neutrophil exudates (Fig. 2). The overall lung tissue structure in mice in groups C and D appeared to be significantly damaged, when compared to that in mice in group A. However, the damage was significantly less than that in group B. The lung tissue structures of mice in these two groups were generally intact (slight destruction of the alveolar structure was observed), only small amounts of neutrophil exudates were observed in the alveolar cavity, there was mild thickening of the alveolar septum (the thickness was 2-4 times that of that in group A), and mild congestion and neutrophil exudates were observed in the interstitial spaces. However, the overall pathological damage in mice in group D was slightly milder, when compared to that in mice in group C (pathological evaluation scores: 0.0625 and 0.0990, respectively). The pathological presentation in group E was similar to that in group D, while the pathological presentation in group F was close to that in group C, but had slightly more severe neutrophil exudation, alveolar septal thickness, and degree of interstitial congestion (the evaluation scores for groups E and F were 0.0781 and 0.1024, respectively). The overall pathological evaluation scores in groups G–J (0.0990, 0.0746, 0.0876 and 0.0793, respectively) were better than those in groups C-F. However, with regard to the four groups, protein deposition in the alveolar cavity was significantly greater in group G than in the other three groups, while interstitial lesions and alveolar septal thickening were greater in group J than in the other three groups. In groups C–J, group D and H had the lowest pathological damage scores (0.0625 and 0.0746, respectively), while groups C and G had the highest scores (both, 0.0990) (Table 3, Fig. 3).

**Figure 2.**
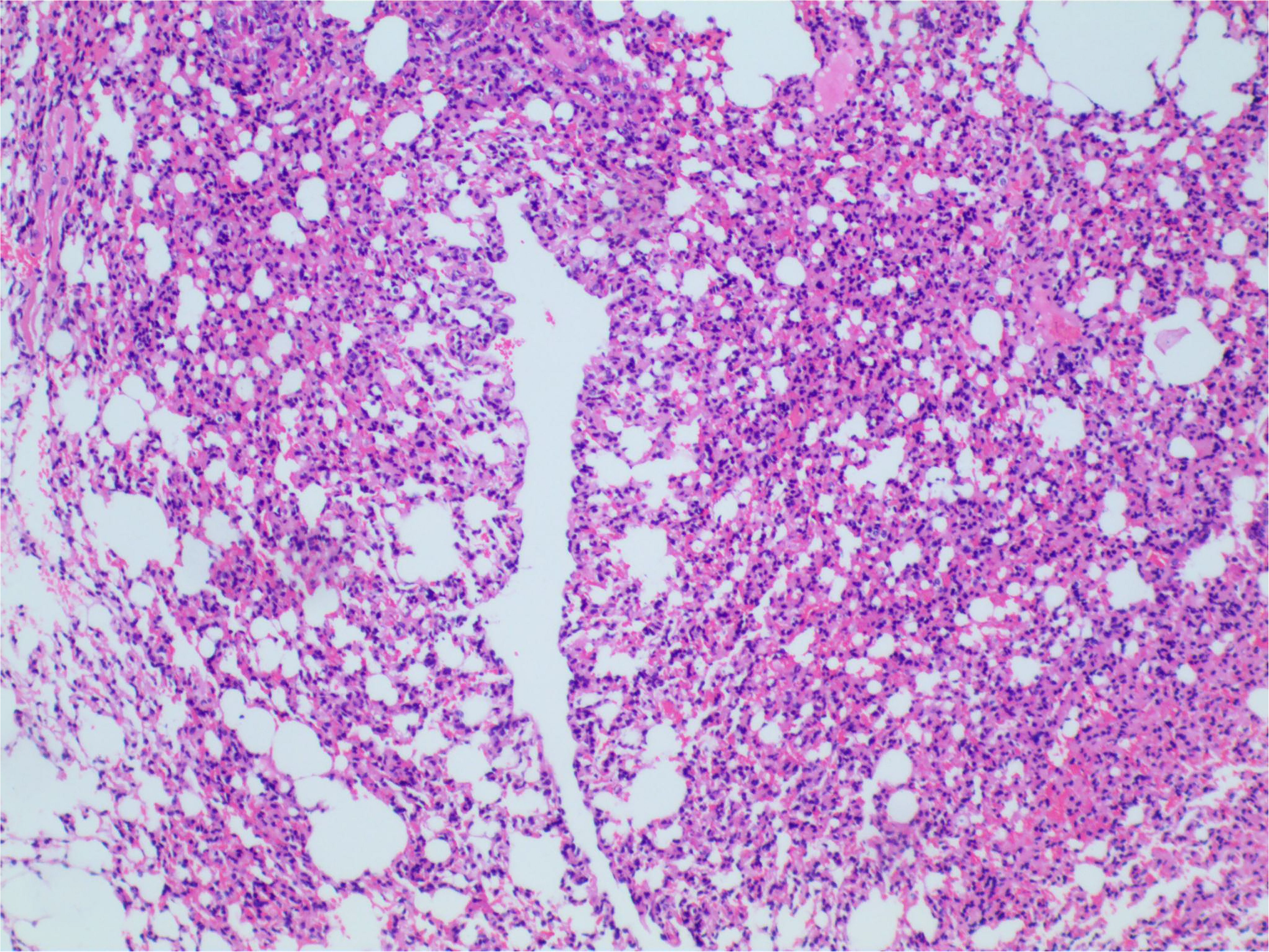
The display of pathological test results in control groups and LPS-only groups. Fig. 2A, 2B and 2C mean control groups which pathological manifestation is present by 100×, 200× and 400× magnified, respectively. The lung tissue structure was clear, the alveolar cavity was clean, the alveolar septum was thin, and there was an absence of interstitial congestion, an absence of neutrophil elevation and migration, and an absence of protein-like substance deposition; Fig. 2D, 2E and 2F mean LPS-only groups which pathological manifestation is present by 100×, 200× and 400× magnified, respectively. It can be seen that the normal structure of the lung tissues had nearly disappeared, the alveolar cavity was significantly reduced or was filled with pink protein-like substances, the alveolar septum was significantly thickened (thickness increased by more than 4 times compared with group A), and there was interstitial congestion and large amounts of neutrophil exudates.

**Figure 3.**
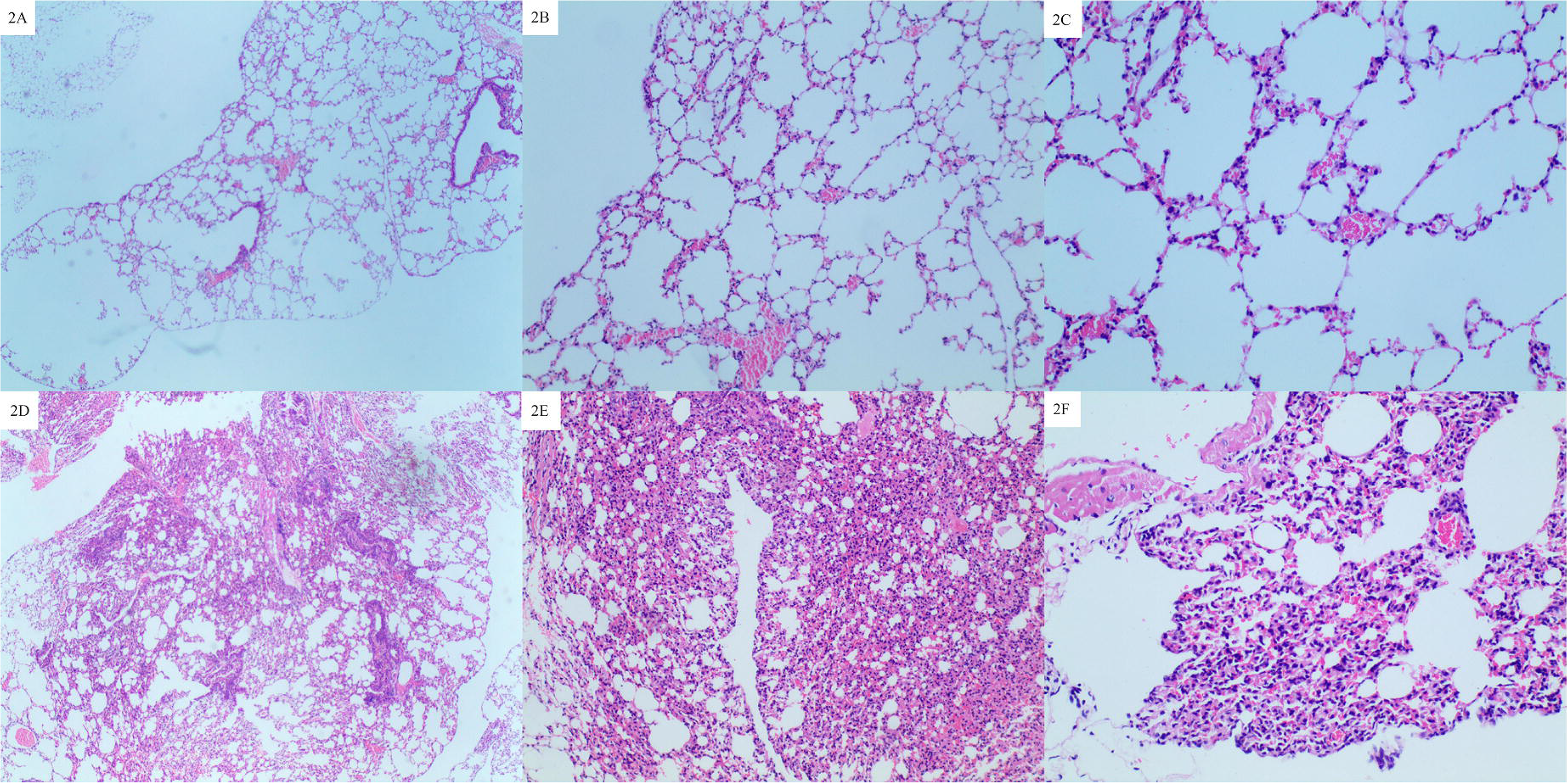
The display of pathological test results among group C~J. Fig. 3A~3H present the pathological test results among group C~J in 400x microscope display, respectively. In group C and group D, the overall lung structure was more obvious damaged than that of group A, but significantly lighter than group B. The structure of the lungs of the two groups remained substantially intact (a small part of the alveolar structure was seen to be destroyed), there was only a very small amount of neutrophils in the alveolar space, and the alveolar septum was slightly thickened (2-4 times the thickness in group A). Mild hyperemia and a small amount of neutrophil exudate were seen. However, overall, pathological lesions in group D were slightly milder than those in group C; pathological findings in group E were similar to those in group D; while those in group F were similar to those in group C, but neutrophil exudation, alveolar septum in alveolar spaces, the thickness and interstitial hyperemia were slightly greater than in group C. The overall pathological score of group G ~ J was better than that of group C ~ F. However, compared with the four groups, the protein deposits in the alveolar space of the G group were significantly more than the other three groups(H ~ J), while the lung interstitial lesions and alveolar septum thickening in the group J were better than the other three groups.

**Table 3.** Pathological evaluation scores for lung tissues in various groups of mice

| Group | Neutrophils<br>in<br>the<br>alveolar<br>cavity | Neutrophils in<br>the interstitial<br>spaces | Protein<br>deposition<br>in<br>air<br>spaces | Alveolar septal<br>thickness | Total<br>score* |
| --- | --- | --- | --- | --- | --- |
| A | 0 | 1 | 0 | 0 | 0.0243 |
| B | 2 | 2 | 2 | 2 | 0.1493 |
| C | 1 | 2 | 1 | 1 | 0.0990 |
| D | 1 | 1 | 0 | 1 | 0.0625 |
| E | 1 | 1 | 1 | 1 | 0.0781 |
| F | 1 | 2 | 1 | 2 | 0.1024 |
| G | 1 | 2 | 0 | 1 | 0.0990 |
| H | 1 | 1 | 1 | 1 | 0.0746 |
| I | 1 | 2 | 1 | 1 | 0.0876 |
| J | 1 | 2 | 1 | 2 | 0.0793 |

\*: Mean evaluation score values for surviving mice in each group

### IL-6 and IL-8 test status

#### IL-6

The IL-6 concentrations of various groups of BALB/c mice (including the blank control group) had significant differences before and after LPS immunization (*P*<0.05). The overall differences in IL-6 concentration after LPS immunization among the groups were statistically significant (*P*<0.05). Among these groups, the LPS only group (group B) had the highest IL-6 concentration, while group E had the lowest IL-6 concentration. Intergroup comparisons revealed that there were no statistically significant differences among groups C, H and B (*P*>0.05). Group E exhibited significant differences when compared with groups B, C and H (*P*<0.005), but no significant difference was found when compared with the other groups (*P*>0.05) (Table 4, Fig. 4A).

**Figure 4.**
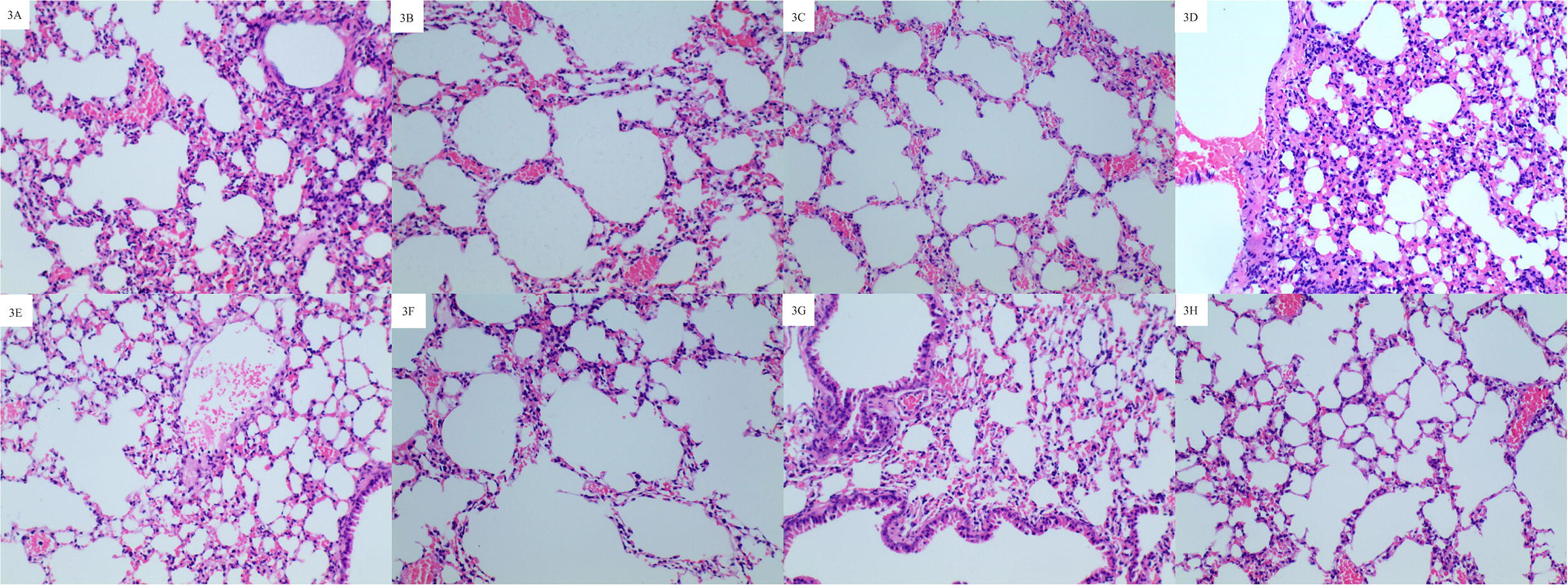
Comparison of IL-6 and IL-8 levels among groups. Fig. 4A: Comparison of IL-6 levels in various groups of mice before and after immunization. Fig. 4B: Comparison of IL-8 levels in various groups of mice before and after immunization.

**Table 4.**
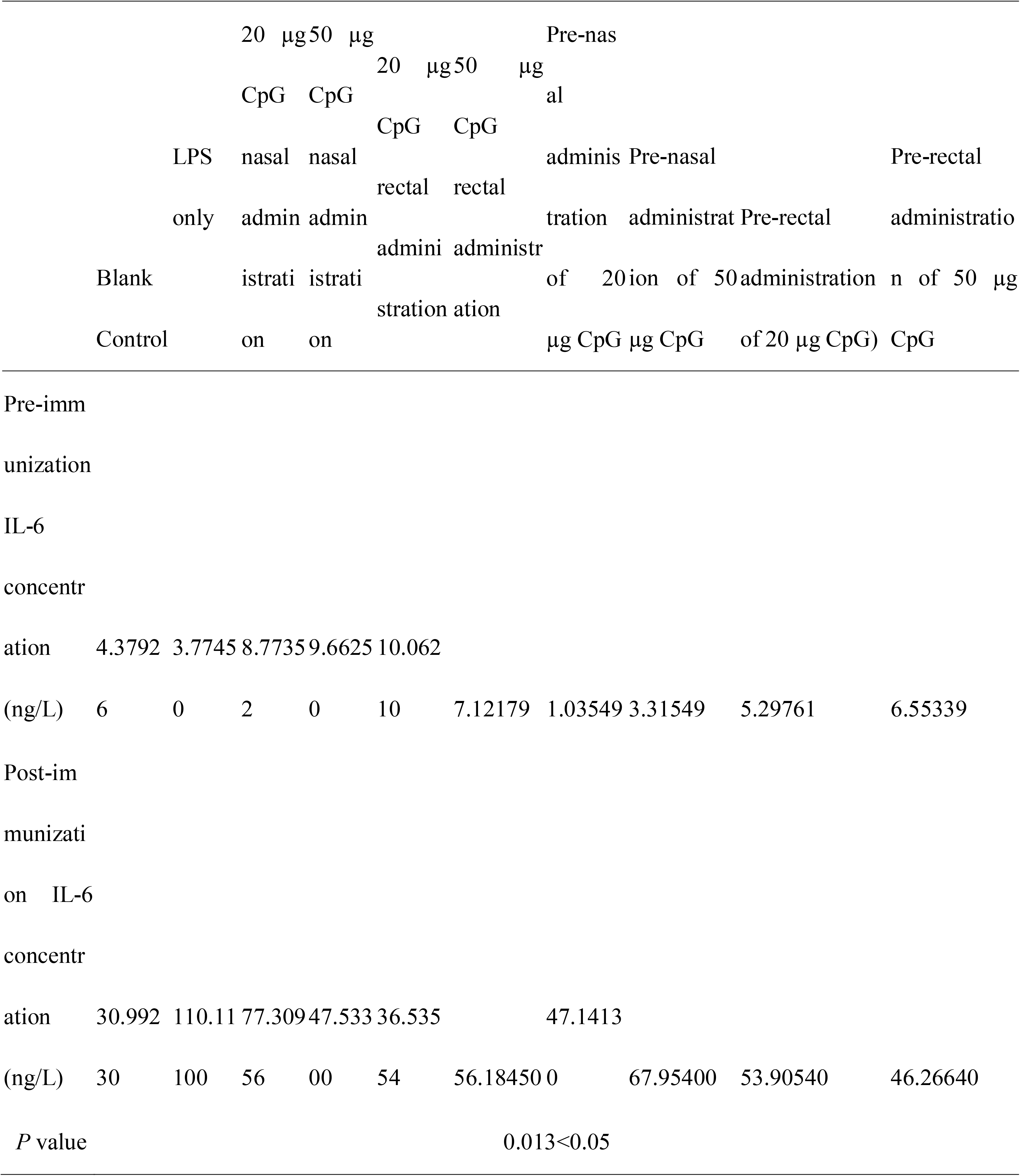
Comparison of IL-6 levels in various groups of mice before and after immunization

#### IL-8

IL-8 concentrations among the groups of BALB/c mice (including the blank control group) had significant differences before and after immunization (*P*<0.05). The overall differences in IL-8 concentration after LPS immunization among the groups were statistically significant (*P*<0.05). Among these groups, the LPS only group and group H (pre-nasal administration of 50 μg of CpG ODN group) had similar IL-8 concentrations, and these two groups had statistically significant differences when compared with group A (blank control group) (*P*<0.05). The comparison of concentrations between the other groups revealed no significant differences (*P*>0.05). In addition, group I (pre-rectal administration of 20 μg of CpG ODN group) had the lowest IL-8 concentration, which was statistically significant when compared with groups B, E and H (*P*<0.05, Table 5, Fig. 4B).

**Table 5.**
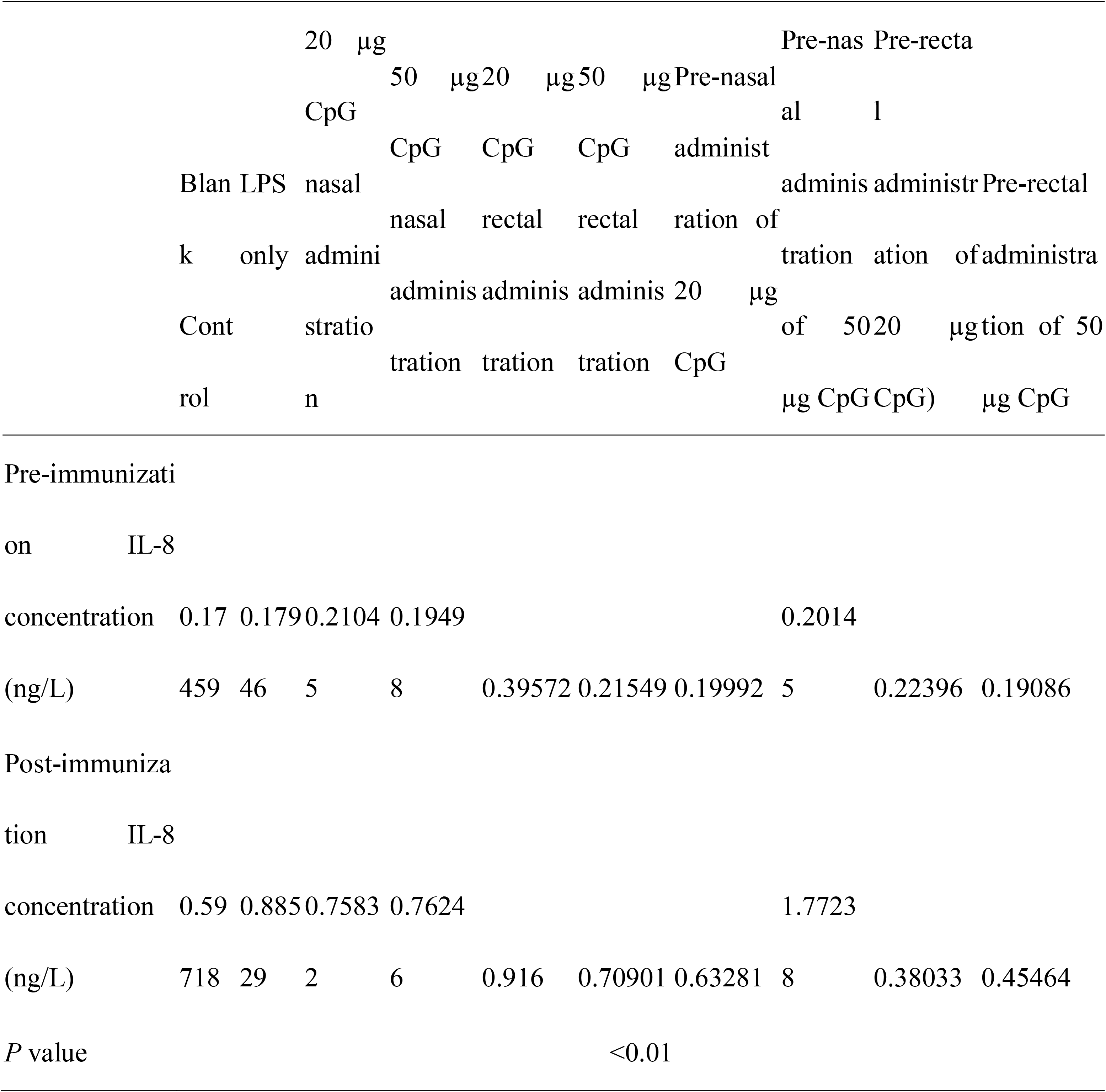
Comparison of IL-8 levels in various groups of mice before and after immunization

## Discussion

Since Ashbaugh (Ashbaugh DG, et al., 1967.) first reported ARDS, the mortality rate of this disease has remained high (Bellani G et al., 2016). One of the reasons for this is that the pathogenesis of ARDS is very complex. Furthermore, it is also due to the aforementioned inflammatory responses that induce the production of autoimmune responses, which result in damage to other non-related organs and makes treatment more challenging. At present, it is clear that (i) inflammatory responses in ARDS induce mitochondrial damage-associated molecular patterns (DAMPs), which activate alveolar macrophages through Toll-like receptor (TLR) and NOD-like receptor (NLR) pathways. The latter pathway results in the release of inflammatory factors, and mobilizes macrophages and neutrophils in blood to aggregate at damage sites. Excessive neutrophils and persistently activated macrophages result in extensive pulmonary epithelial and endothelial cell damage, which in turn causes damage to the alveolar-capillary barrier. The destruction of this barrier enables protein-rich edema fluid to enter the alveoli, resulting in alveolar fluid accumulation (alveolar edema), thereby affecting normal gaseous exchange. (ii) Ubiquitination plays an important role in regulating key proteins that cause ARDS, since this process can induce the secretion of cytokines (mainly, IL-6, IL-8, IL-1β and TNF), decrease pulmonary surfactant levels, and reduce ion channel function (including Na+ /K+ -ATPase channels, and epithelial sodium channel). (iii) The consumption of alveolar surfactants and increased NETosis by neutrophils further worsens pulmonary cellular functions. (iv) DAMPs can also directly cause an increase in vascular permeability without relying on granulocytes, resulting in a repeated cycle of inflammation (Han S and Mallampalli RK, 2015). Since these processes involve multiple immune responses and inflammatory mediators, treatment is difficult. Existing methods are mainly directed against the primary disease, and merely few methods are directed against lesions in lung tissues. Limiting or eliminating these responses to decrease the severity of lung tissue injury and shortening the duration of inflammatory responses to prevent the occurrence or development of ARDS requires the consideration of whether mucosal immunity can play a role in limiting or eliminating the occurrence or development of ARDS.

The mucosa is an important physiological barrier in the innate immune system, which prevents invasion by pathogenic microorganisms. It constitutes as a physiological barrier and immune protection, since the mucosa itself has its own immune system, termed mucosal immunity. The production of mucosal immunity mainly involves M cells, DCs, T lymphocytes and B lymphocytes. In addition, the aforementioned sensitized B lymphocytes can migrate out of the mucosal sites, in which these are sensitized to pass through lymphatic vessels, the thoracic duct and blood, before returning to its initial sensitization site (this process is termed, lymphocyte homing). This process induces mucosal immunity at multiple distal sites. Systemic immunity is also activated by this process, but the effect is relatively weak. Therefore, activated B-cells, IgA, T-cells and DCs all constitute the mucosal defense system, and play roles in neutralizing and eliminating pathogens to promote mucosal repair.

Recent studies (Chen J, et al., 2016; Li X, et al., 2016) have shown that adjuvant CpG ODN is a ligand for TLR9, and is a good stimulant for mucosal immunity, since it can induce mucosal immunity alone. When CpG ODN is engulfed by mucosal cells, it interacts with TLR9 in the cytoplasm to form a complex that activates immune responses through a myeloid differentiation factor 88 (MyD88)-dependent pathway. MyD88 and TNF receptor associated factor 6 (TRAF6) can activate NF-kappa-B (NF-κB)-inducing kinase, which subsequently activates the inhibitor of NF-κB (IκB) kinase (IKK). This causes the phosphorylation of IκB and NF-κB to be released into the cell nucleus and activate target genes, which activates the gene transcription of relevant cytokines. CpG ODN can also induce the transcription and expression of high levels of granulocyte colony-stimulating factor (G-CSF) (Nardini E, et al., 2005), tumor necrosis factor-alpha (TNF-α), IL-6 (Zhu FG and Marshall JS, 2001) and other cytokines through other cell signaling pathways.

The intratracheal instillation of LPS into BALB/c mice was used to successfully generate the ARDS model before analyzing the protective effects of the intranasal and transrectal induction of mucosal immunity. From these pathological results, it can be observed that the intranasal administration of 50 μg of CpG ODN after LPS immunization in mice resulted in the least severe lung tissue injury. Furthermore, IL-6 and IL-8 concentrations were lower, which was second to mice treated with the rectal administration of 20 μg of CpG ODN. From the survival outcomes of various groups, it can be observed that mice in the nasal administration of 20 μg of CpG ODN after LPS immunization group, rectal administration of 20 μg of CpG ODN after LPS immunization group, and pre-nasal administration of 20 μg of CpG ODN group all had 100% survival rates. Furthermore, mice in the blank control group, LPS only group, nasal administration of 50 ug of CpG ODN after LPS immunization group, rectal administration of 50 ug of CpG ODN after LPS immunization group, and pre-rectal administration of 50 μg of CpG ODN group all had 80% survival rates, while the survival rate of the other groups was only 60%. Therefore, even though there were intergroup differences in IL-6 and IL-8, mice in the group where 50 μg of CpG ODN was intranasally administered after LPS immunization had the most minor pathological damage and relatively good survival outcomes. In contrast, the nasal and rectal administration of CpG ODN in BALB/c mice before LPS immunization did not appear to exhibit any significant protective effect. However, it is worth noting that regardless of the timing of CpG ODN administration, the IL-6 and IL-8 concentrations are relatively lower when CpG ODN was intrarectally administered. This may explain how mucosal immunity can correct excessive inflammatory responses in ARDS (Reiss LK, et al., 2012), enabling the regulation of the imbalance between pro-inflammatory and anti-inflammatory responses, thereby alleviating tissue injury. This causes the levels of inflammatory mediators to decrease, which can reduce tissue damage and accelerate repair.

In conclusion, the present study demonstrates that CpG ODN has protective effects for ARDS in mice, which may be due to the stimulation of CpG ODN on the mucosa, inducing the production of large of IgA to neutralize various microorganisms, which in turn reduces inflammatory cytokines, and leads to less pathologic injury. Furthermore experiments are needed with more mice, in order to determine the protective effect of CpG ODN to exogenous ARDS.

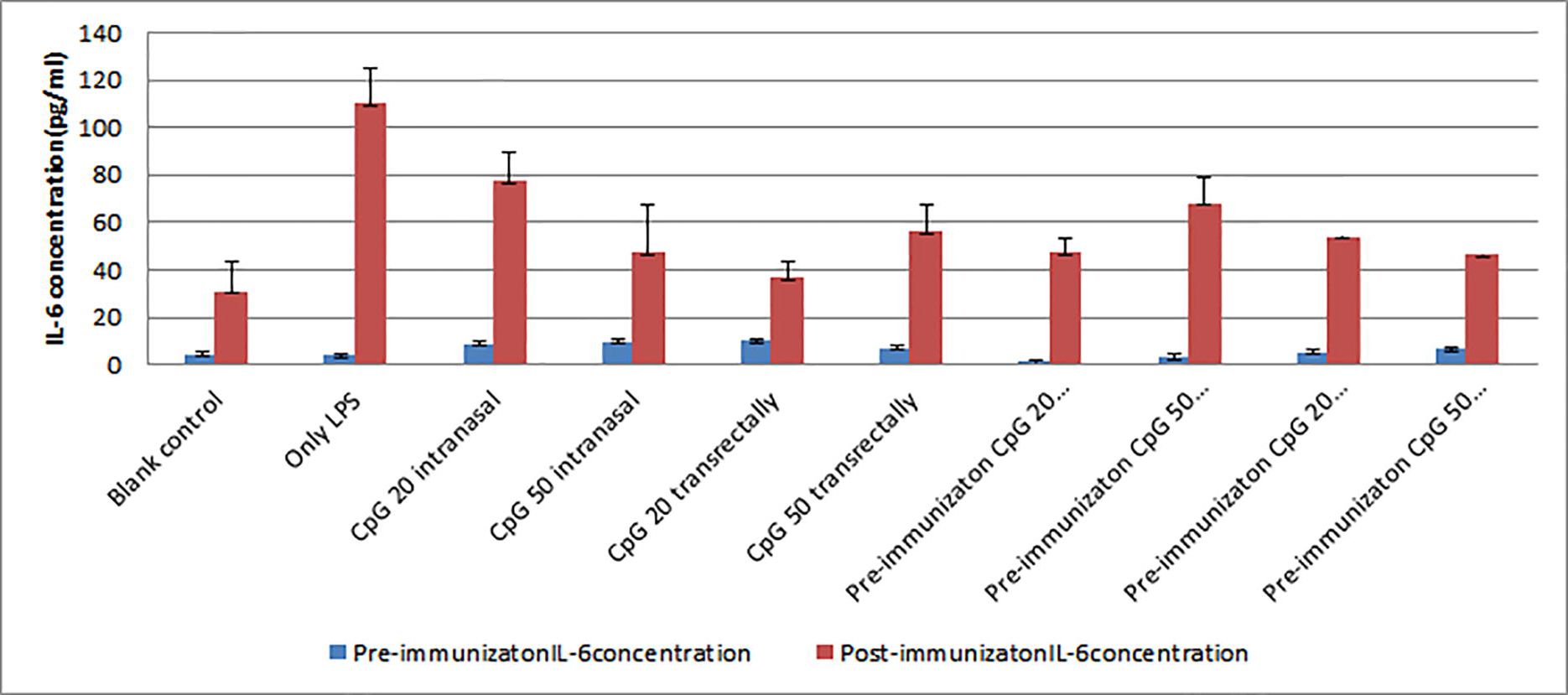

## Acknowledgements

This study was supported by Science and Technology Committee of Tongzhou District, Beijing City (grant number: KJ2016CX009-08).

